# The regulatory effect of Ca2+/calmodulin dependent protein kinase II mediated by receptor interacting protein kinase 3 on necroptosis of hypertrophic cardiomyocytes

**DOI:** 10.1101/2025.09.29.679355

**Authors:** Jingjing Zhang, Huijie Yu, Yuxin Yang, Ajie Liu, Peng Wang

## Abstract

Receptor interacting protein kinase 3 (RIPK3) plays a crucial role in the signaling pathway of necroptosis and calcium/calmodulin dependent protein kinase II (CaMKII) is a novel substrate for RIPK3 induced necroptosis. The aim of this study is to investigate the regulation and mechanism of RIPK3 on AngII induced cardiomyocyte hypertrophy.AngII was used to stimulate myocardial cells for 72 hours, inducingcardiomyocytehypertrophy; Use RIPK3 inhibitor GSK’872 to reduce RIPK3 expression.Detect indicators related to myocardial hypertrophy, cell damage, necroptosis,CaMKII activation and gene expression, oxidative stress, mitochondrial membrane potential, etc. After AngII stimulation of cardiomyocytes, the expression of hypertrophy markers ANP and BNP increased, LDH release increased, ATP levels decreased, splicing factors ASF and SC35 expression increased, CaMKII oxidation and phosphorylation levels increased, and CaMKIIδ alternative splicing was disrupted. However, treatment with GSK’872 can alleviate myocardial dysfunction, inhibit CaMKII activation, correct CaMKIIδ variant splicing disorder and ultimately alleviating myocardial hypertrophy. In addition, pretreatment with RIPK3 can reduce the accumulation of reactive oxygen species (ROS) induced by AngII, decrease the activity of ASF and SC35, and restore mitochondrial membrane potential. RIPK3 inhibitor GSK’872 can inhibit the activation of CaMKII, alleviate necroptosis and oxidative stress to alleviate myocardial hypertrophy. It has a protective effect on myocardial hypertrophy and is expected to become a new targeted drug for clinical treatment of dilated cardiomyopathy and heart failure.

---

Cardiac hypertrophy is an adaptive response of myocardial cells to various stimuli^[1]^. It is a compensatory response of the body to stress or stimulation, which can lead to arrhythmia and heart failure. Although there are various molecular mechanisms involved, it is still difficult to treat. Early manifestations include increased myocardial cell volume, increased protein synthesis, and enhanced myocardial contractility^[2]^. Cardiac hypertrophy can be divided into physiological hypertrophy and pathological hypertrophy. Physiological hypertrophy occurs during exercise, development, and pregnancy, and maintains cardiovascular function over time. Pathological myocardial hypertrophy refers to the structural and functional changes in myocardial cells caused by pressure and/or volume overload, as well as excessive stimulation of various growth factors and/or hormones under pathological conditions. Persistent pathological myocardial hypertrophy can cause dilated cardiomyopathy, ultimately leading to heart failure. Therefore, it is very important to search for new therapeutic targets for myocardial hypertrophy^[3]^.

Apoptosis is a strictly controlled active process with a well-established regulatory mechanism, while necrosis is considered an accidental or passive cell death^[4]^. There are many morphological changes in apoptosis, including cell shrinkage, pseudopodia disappearance, chromatin condensation, nuclear membrane shrinkage, nucleolar division, and ultimately the formation of apoptotic bodies^[5]^. Recent research progress has revealed a new type of necrosis called regulatory cell death, also known as necroptosis, which is morphologically distinct from apoptosis and necrosis. It exhibits morphological features of necrotic cells induced by plasma membrane rupture and contents leakage^[6-7]^. Inhibiting necroptosis has a protective effect on aging induced ischemic hearts^[8]^; Necroptosis inhibitor has beneficial effects on cell viability and cardiac function in a chronic myocardial hypertrophy model^[9]^. However, it is still unclear whether necroptosis plays a key role in myocardial hypertrophy.

Several studies have shown that receptor interacting protein kinase 3 (RIPK3) is a key regulatory factor in the necroptosis signaling pathway^[10]^. RIPK3 can bind to receptor interacting protein kinase 1 (RIPK1) through the RIP homotypic interaction motif (RHIM) to form necrosomes, triggeringnecroptosis^[11]^. Studies have shown that upregulation of RIPK3 plays a critical role in ischemic and oxidative stress-induced myocardial cell necroptosis, leading to myocardial remodeling and heart failure in addition to apoptosis and inflammation. Upregulation of RIPK3 induces necroptosisof myocardial cells by activating CaMKII. Targeting the RIPK3-CaMKII pathway can protect the heart from the effects of ischemic and oxidative stress-induced necrotic cell death, myocardial remodeling, and heart failure^[4]^. However, the expression and role of RIPK3 in myocardial hypertrophy are still unclear.

CaMKII is a multifunctional protein kinase that regulates cell biology and plays an important physiological role in the heart. It regulates the excitation contraction and excitation transcription coupling of myocardial cells by acting on ion channels^[12]^, and can maintain normal diastolic function of the heart and stable Ca^2+^concentration in myocardial cells. However, sustained activation of CaMKII has been recognized as a central mediator of programmed cell death in cardiovascular diseases (including necroptosis), which is detrimental to cardiac integrity and function. In addition, its activation mediates physiological or pathological responses and remodeling under cardiac stress. It has been fully established that CaMKII plays an important role in myocardial hypertrophy, pressure overload induced cardiac hypertrophy and fibrosis, ischemia/reperfusion (I/R) injury, heart failure (HF), post myocardial infarction (MI) remodeling, and ventricular arrhythmia^s[13-18]^. Recently, some studies have revealed the potential role of CaMKII in myocardial hypertrophy. Therefore, identifying the mechanism of CaMKII can help provide new pharmacological targets for myocardial hypertrophy^[19]^.

The excessive activation of a bioactive substance, angiotensin II (AngII), is associated with the occurrence and development of myocardial hypertrophy, and it plays an important role in myocardial hypertrophy and remodeling. AngII induces physiological changes such as vasoconstriction, fibrosis, and cell death, accompanied by mitochondrial dysfunction^[20]^. According to clinical trial analysis, angiotensin-converting enzyme inhibitors are more effective in reducing myocardial hypertrophy than other antihypertensive drugs, indicating that the reversal of myocardial hypertrophy is more significantly affected by changes in angiotensin II than by blood pressure reduction^[21]^.The alternative splicing factors ASF and SC35 are key factors regulating the alternative splicing of CaMKIIδmRNA precursor CaMKIIδ^[22-23]^. Research has found that regulating bispecific tyrosine phosphorylation kinase 1A (Dyrk1A) can induce phosphorylation of splicing factors, causing ASF/SC35 translocation and losing control of alternative splicing regulation^[24]^. Previous studies have found that CaMKIIδinhibitors alleviate myocardial cell hypertrophy induced by angiotensinII, phenylephrine, and electric field stimulation, while overexpression of CaMKIIδB or CaMKIIδC enhances myocardial cell hypertrophy^[25-26]^;ASF binds to protein phosphatase1γ (PP1γ), which can increase alternative splicing of CaMKIIδ and promote the expression of CaMKIIδB/C mRNA variants^27^; Overexpression of myocardial PP1γ enhances CaMKIIδ activity and reduces myocardial hypertrophy^[28]^.

In summary, we investigated the molecular mechanism of necroptosis in the development of myocardial hypertrophy and demonstrated the regulatory effect of RIPK3 inhibitor GSK’872 on CaMKIIδalternative splicing and CaMKII activity to improve AngII induced myocardial cell hypertrophy.

## Results

### Angiotensin II stimulates cultured myocardial cells in vitro, inducing myocardial cell hypertrophy. The expression of hypertrophy markers and RIPK3 in myocardial cells is upregulated; The activity of CaMKII δ in cardiomyocytes is increased, and the alternative splicing disorder of CaMKII δ leads to an increase in necroptosis of cells

After stimulation with angiotensin II, the protein expression of myocardial hypertrophy markers, atrial natriuretic peptide, and brain natriuretic peptide (ANP and BNP) in cultured cardiomyocytes was significantly increased, indicatingthesuccessful establishment of a myocardial hypertrophy model in this study. Western blot analysis showed that compared with the control group, the protein expression of RIPK3 and p-RIPK3 in the AngII group cardiomyocytes was significantly increased (P<0.05). Therefore, myocardial cell hypertrophy greatly enhances the expression of RIPK3 and phosphorylated RIPK3 (Fig.1). Subsequently, the oxidation and phosphorylation levels of CaMKII in hypertrophic cardiomyocytes also significantly increased (P<0.05). Chronic activation of CaMKII leads to cellular remodeling, ultimately resulting in myocardial cell damage caused by changes in Ca^2+^processing, ion channels, intercellular coupling, and metabolism. My study used quantitative real-time PCR to detect the expression of CaMKII δ A, CaMKII δ B, and CaMKII δ C mRNA. It was found that in the AngII stimulated myocardial cell group, CaMKII δ A and CaMKII δ B decreased, while CaMKII δ C increased, indicating a disruption in CaMKII δ alternative splicing after myocardial cell hypertrophy (Fig.2). Through TUNEL staining, the results showed that the proportion of positive myocardial cells increased after AngII stimulation, indicating an increase in the level of myocardial cell apoptosis (P<0.05). The Western Blot test results showed that the expression level of cleaved caspase 3 also increased synchronously after AngII pretreatment (P<0.05). In addition, the results of LDH release and ATP activity detection in myocardial cell culture medium showed that the release of LDH was significantly increased (P<0.05) and ATP content was significantly decreased (P<0.05) in the AngII stimulated myocardial cell group, indicating that AngII has a damaging effect on the organelles of cultured myocardial cells (Fig.3). The above data indicate that after myocardial cell hypertrophy, myocardial damage occurs, CaMKIIδ activity increases, CaMKIIδ alternative splicing disorder occurs, and necroptosis intensifies.

**Fig. 1.**
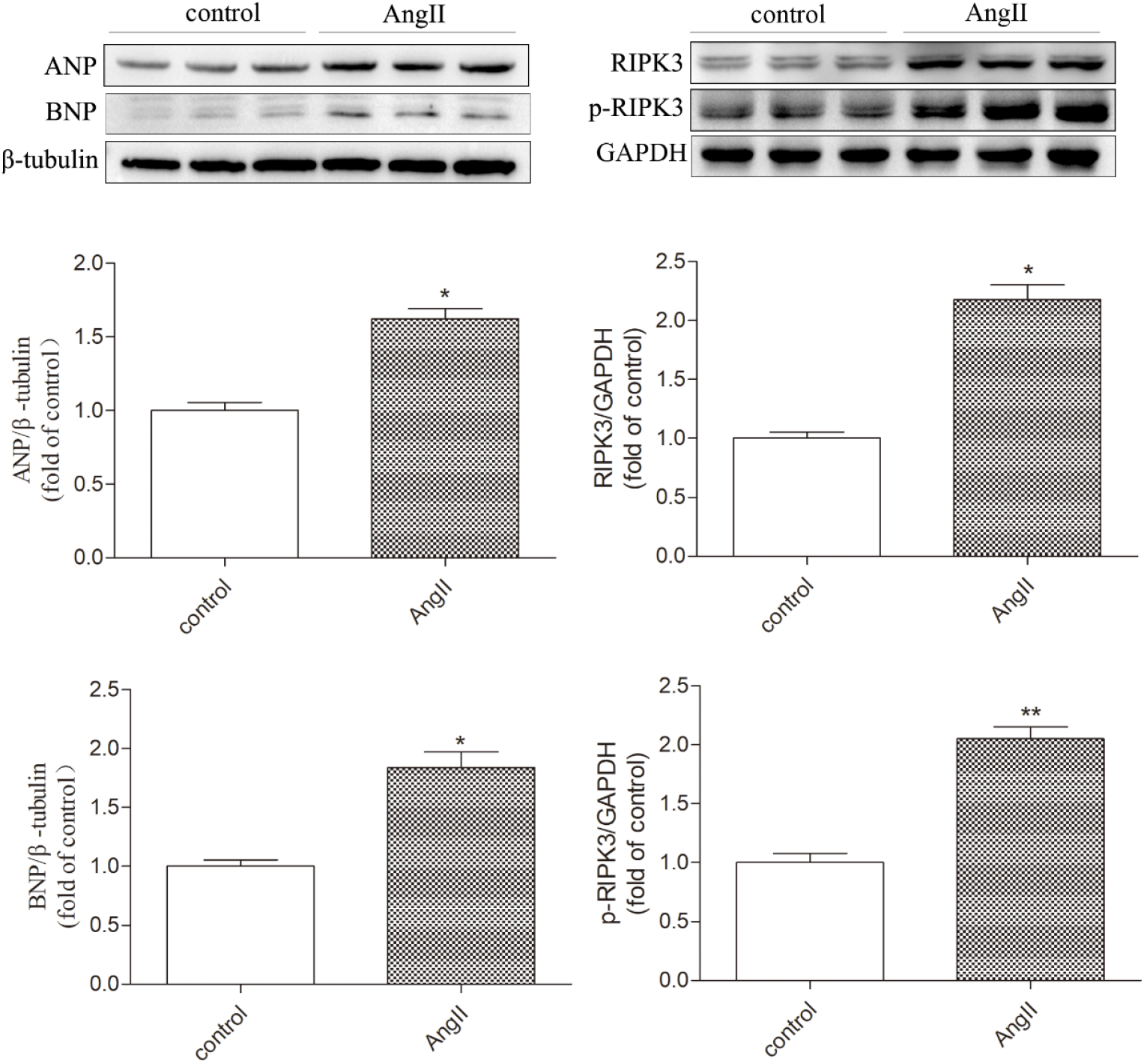
AngII induces myocardial cell hypertrophy, with increased expression of RIPK3 and its phosphorylation. Cultivate primary cardiomyocytes with medium containing 100 μ M AngII or without AngII for 72 hours, and detect the protein expression levels of ANP, BNP, RIPK3, and p-RIPK3 in cultured cardiomyocytes by Western Blot. The data is the mean ± SEM, compared with the control group * P<0.05, **P<0.01,n=6.

**Fig. 2.**
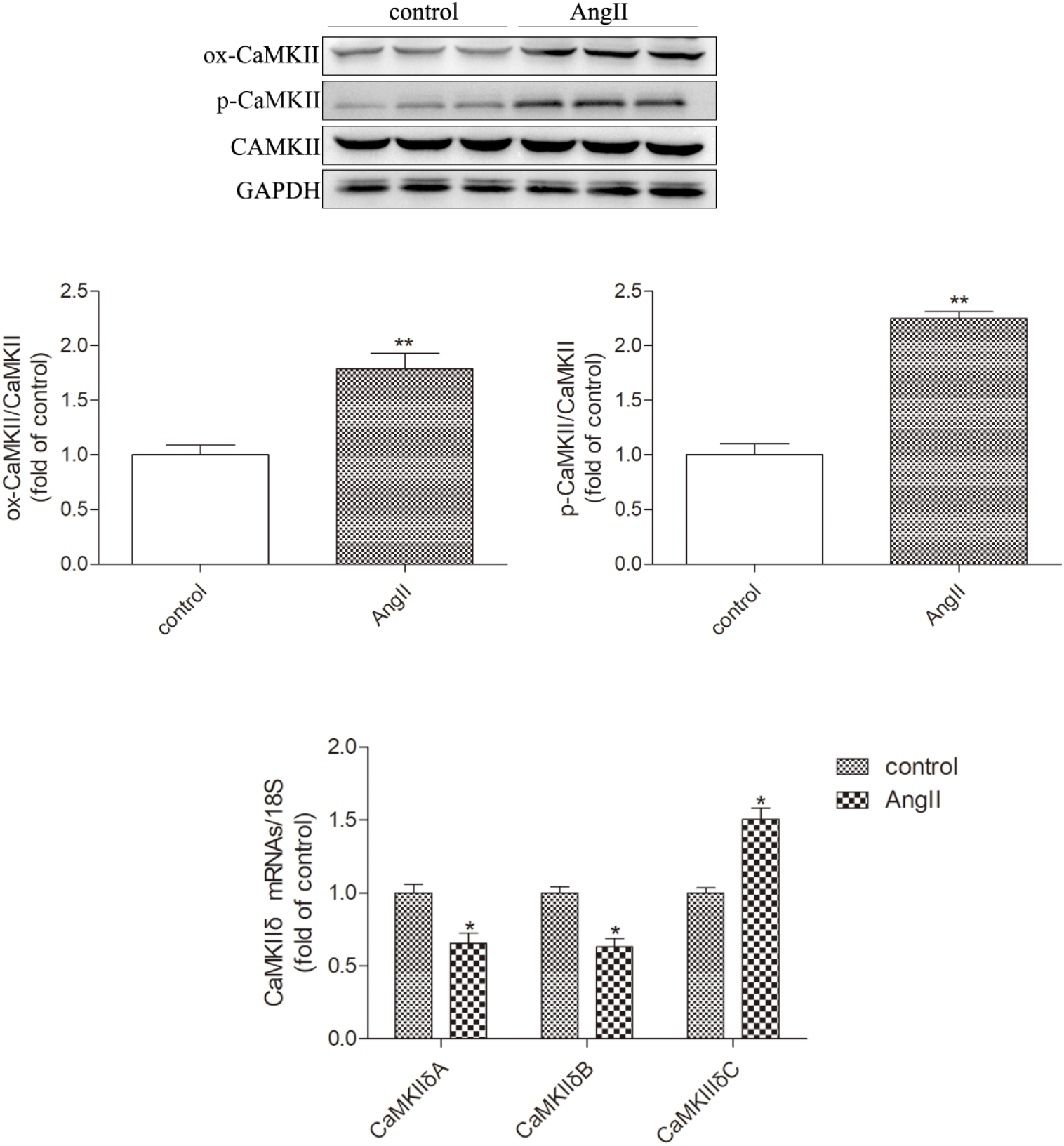
AngII induces CaMKII activation and necrotic apoptosis in cardiomyocytes. Western blot was used to detect the expression levels of CaMKII oxidation (ox CaMKII), CaMKII phosphorylation (p-CaMKII), and total CaMKII. Real time quantitative PCR was used to detect the mRNA expression levels of CaMKII δ A, CaMKII δ B, and CaMKII δ C in myocardial cells. The data is the mean ± SEM. Compared with the control group* P<0.05, **P<0.01,n=6.

**Fig. 3.**
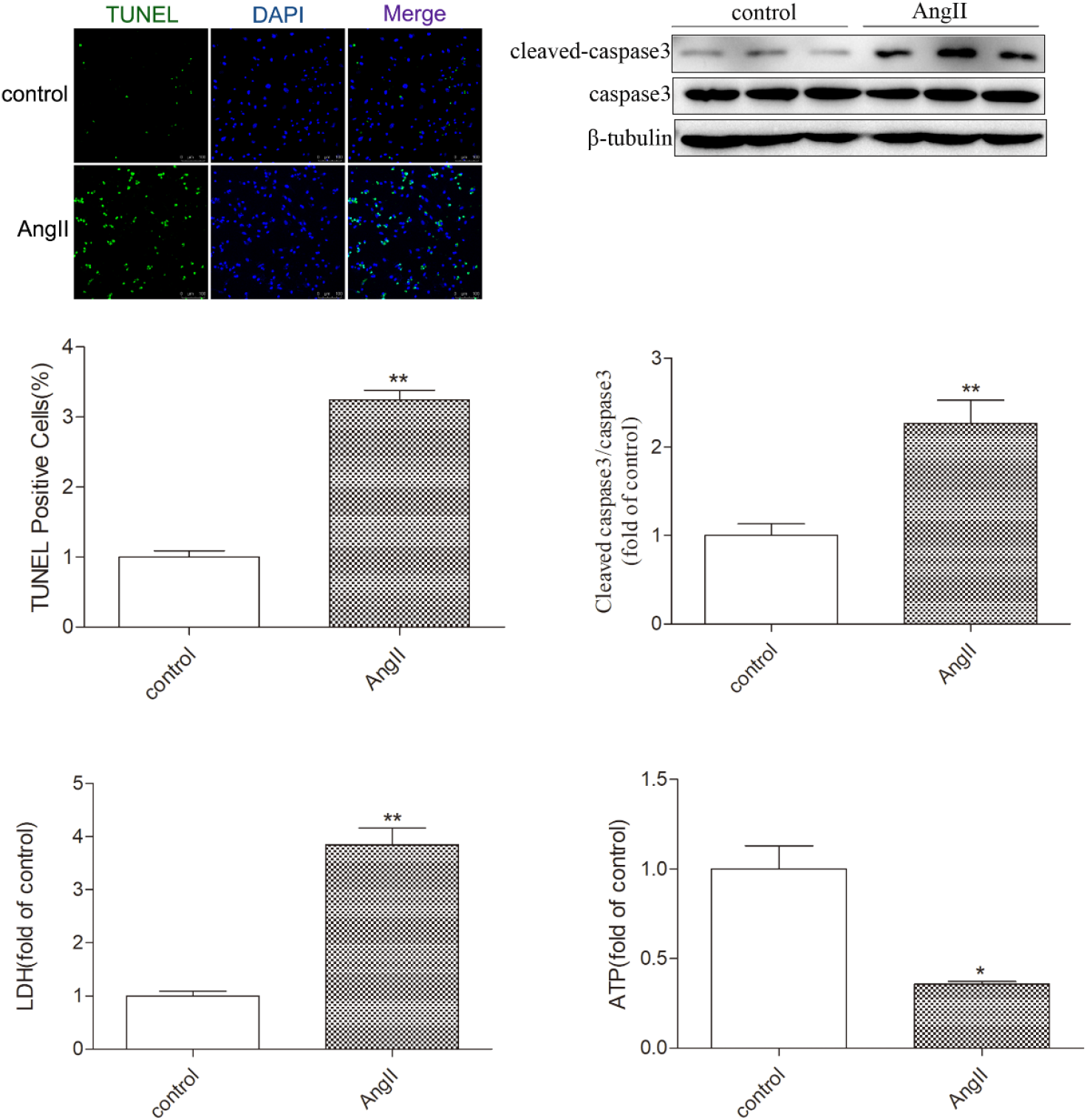
TUNEL staining for detecting myocardial cell apoptosis (Bar=50 μ m). Western Blot was used to detect the expression levels of cleaned-caspase3 and caspase 3 proteins to evaluate the necrotic apoptosis of myocardial cells. Detect the release of LDH and ATP levels in myocardial cell culture medium. The data is the mean ± SEM. Compared with the control group* P<0.05,**P<0.01, n=6.

### GSK’872 inhibits the expression of AngII induced cardiac hypertrophy markers, RIPK3, and p-RIPK3

To investigate the role of RIPK3 in myocardial cell hypertrophy, RIPK3 inhibitor GSK’872 was used to downregulate RIPK3 expression in myocardial cells. Detect the protein expression of ANP and BNP through WB; The results showed that compared with the AngII myocardial cell stimulation group, pre-treatment with GSK’872 significantly inhibited the expression levels of ANP and BNP under AngII stimulation (P<0.05), further demonstrating the inhibitory effect of GSK’872 on AngII induced myocardial hypertrophy in vitro. In addition, WB results showed that the expression level of RIPK3 in the GSK’872 group was significantly lower than that in the control group (P<0.05), and the expression level in the GSK’872+AngII group was also significantly lower than that in the AngII group (P<0.05). The changes in protein expression levels of p-RIPK3 are basically consistent with RIPK3 (Fig.4). Therefore, GSK’872 can effectively inhibit the expression and phosphorylation of RIPK3 in cardiomyocytes after AngII stimulation. In summary, these data indicate that the absence of RIPK3 can effectively improve AngII induced cardiomyocyte hypertrophy.

**Fig. 4.**
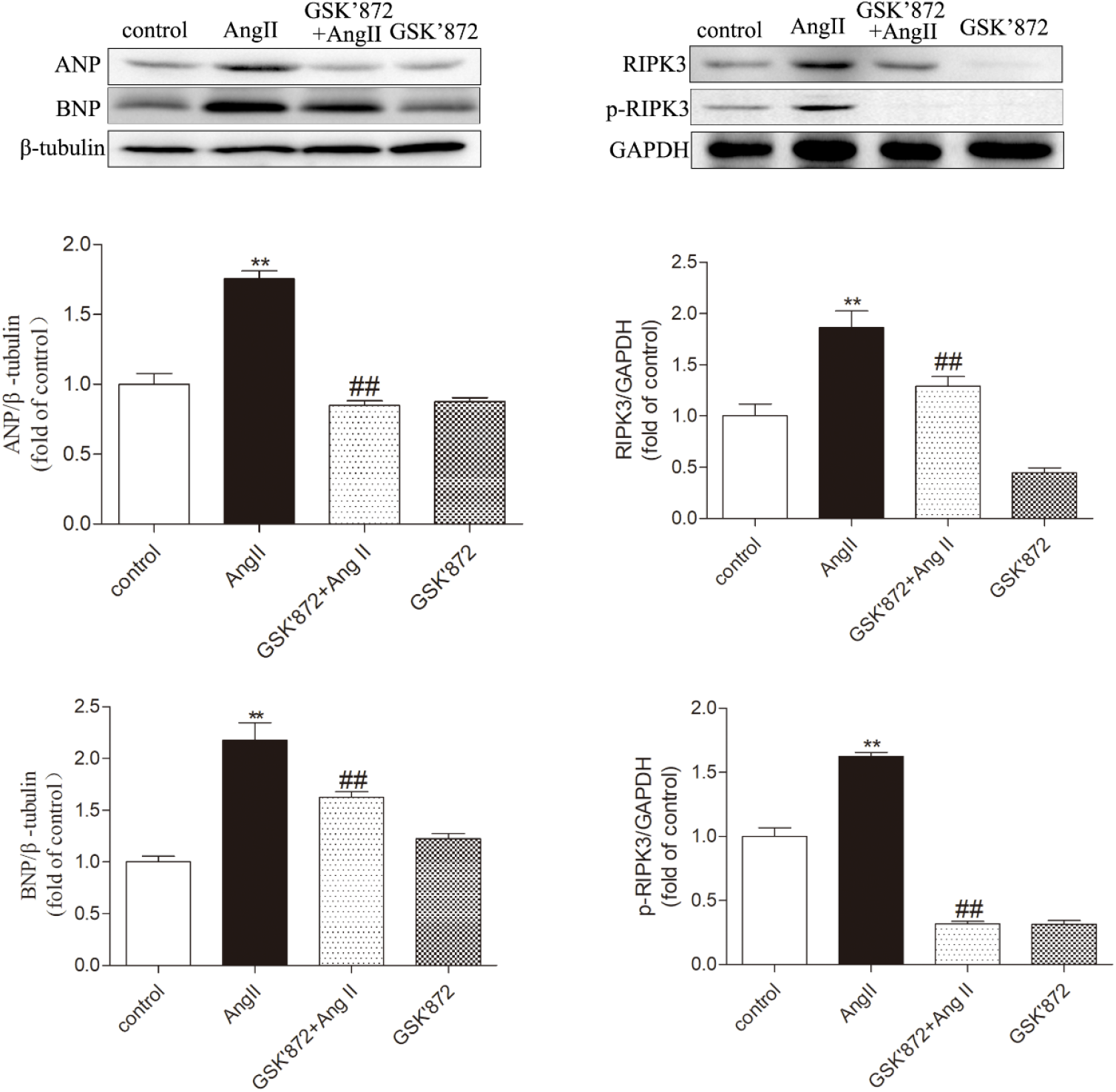
Pretreatment with RIPK3 inhibitor GSK’872 inhibits AngII induced myocardial cell hypertrophy and RIPK3 expression. Primary cardiomyocytes were pretreated or not pretreated with 10 μ M GSK’872, and after 4 hours, they were replaced with culture medium containing AngII (100 μ M) or without AngII. Observe the protein expression changes of ANP and BNP through Western blot analysis to evaluate the degree of myocardial cell hypertrophy. Quantify the protein expression of RIPK3 and p-RIPK3 through Western Blot. The data is represented as mean ± SEM. Compared with the control group** P<0.01, Compared with the GSK’872+AngII group ## P < 0.01,n=6.

### GSK’872 inhibits AngII induced myocardial cell dysfunction, CaMKII oxidation and phosphorylation

Persistent CaMKII activation plays a crucial role in the progression of myocardial hypertrophy to heart failure. Next, we investigated whether RIPK3 regulates myocardial cell hypertrophy by activating CaMKII. Our study showed that in cultured cardiomyocytes treated with GSK’872 and AngII, the overall expression of CaMKII was not different from other groups of cells, but the expression of ox CaMKII and p-CaMKII was significantly decreased compared to cardiomyocytes treated with AngII alone (P<0.05). Therefore, GSK’872 can effectively alleviate AngII induced oxidation and phosphorylation of CaMKII in cardiomyocytes (Fig.5).

**Fig. 5.**
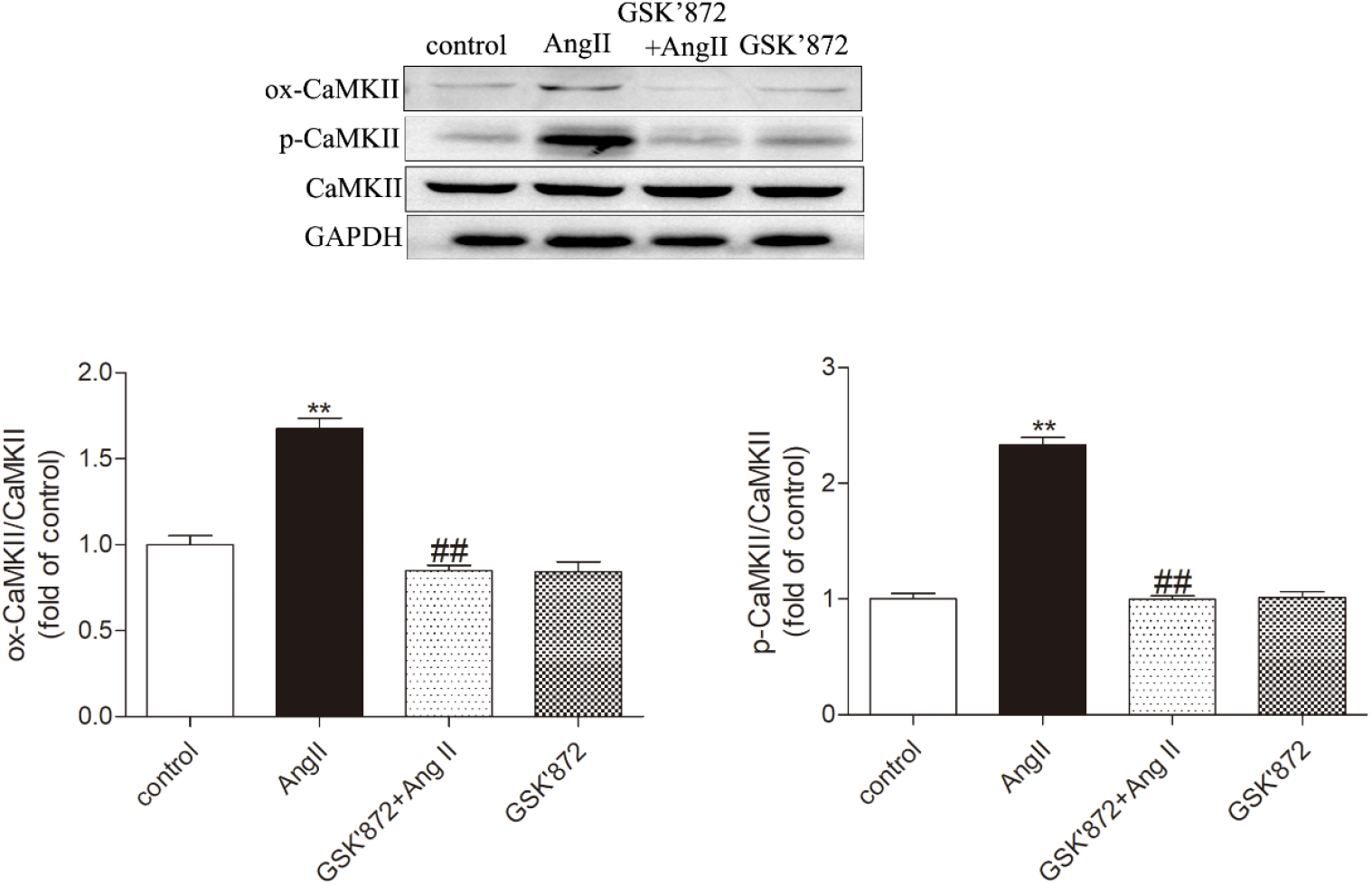
AngII induced myocardial injury and CaMKII activation were relieved after GSK’872 pretreatment. Western blot was used to detect the expression of ox CaMKII, p-CaMKII, and total CaMKII. The data is the mean ± SEM. Compared with the control group* P<0.05,**P<0.01; Compared with the GSK’872+AngII group ^#^ P < 0.05,^##^P < 0.01,n=6.

### GSK’872 inhibits AngII induced expression of ASF and SC35 splicing factors in cardiomyocytes, alleviates CaMKII δ alternative splicing disorder

Our results showed that in cardiomyocytes treated with GSK’872 and AngII, GSK’872 effectively reduced the protein expression of ASF and SC35 induced by AngII in cardiomyocytes; Through GSK’872 targeted therapy, the mRNA levels of CaMKII δ A and CaMKII δ B in cardiomyocytes were upregulated and CaMKII δ C was decreased after AngII pretreatment, thereby alleviating alternative splicing disorder. in summary, Downregulation of RIPK3 by GSK’872 can inhibit the expression of splicing factors in cardiac mast cells and disrupt the expression of CaMKII δ variants (Fig.6).

**Fig. 6.**
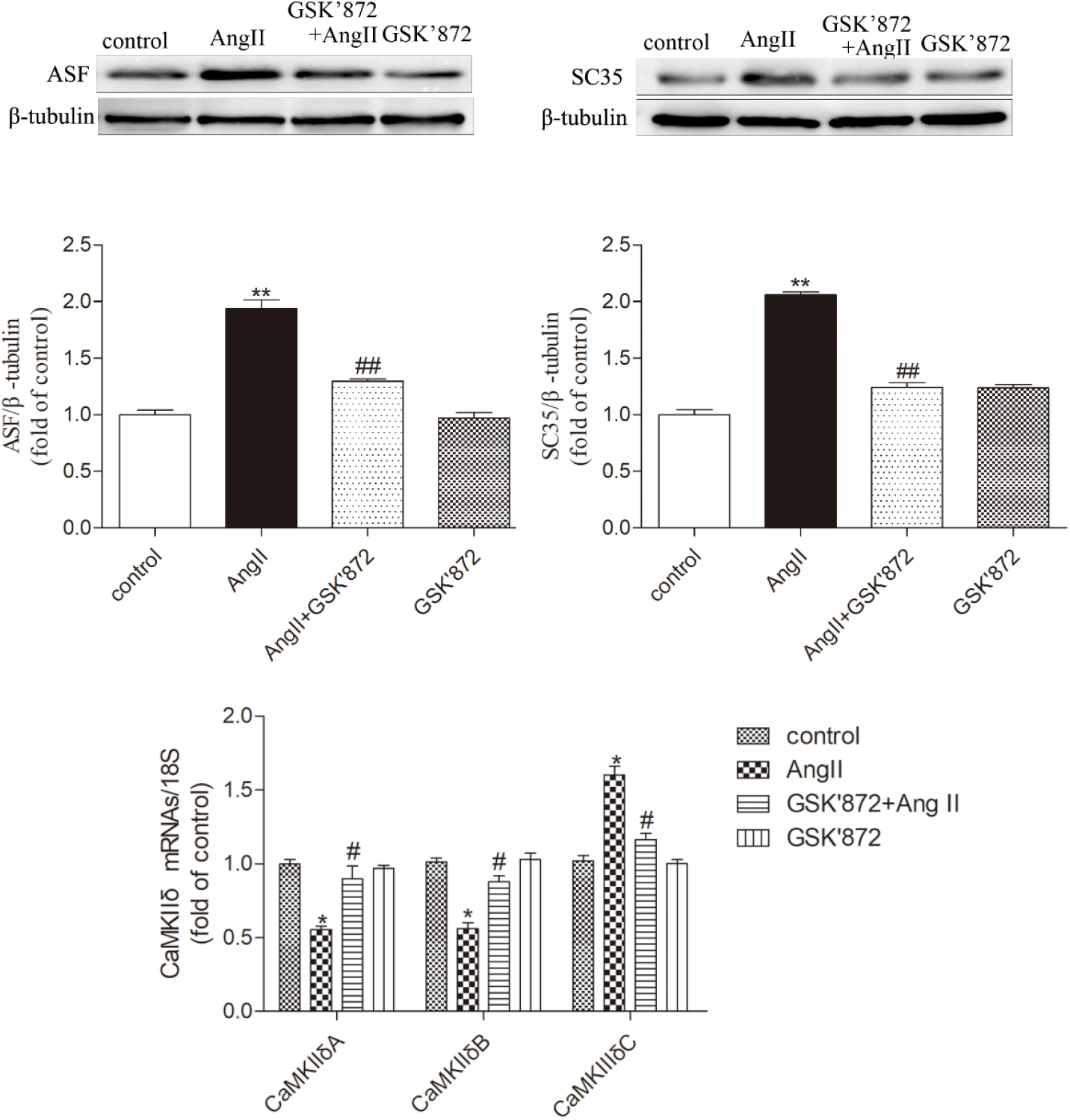
RIPK3 inhibitor 10 μ M GSK’872 pretreatment of cardiomyocytes, Western Blot was used to detect the expression of ASF and SC35 proteins in cardiomyocytes. Using qRT PCR to detect the mRNA levels of CaMKII δ A, CaMKII δ B, and CaMKII δ C in cardiomyocytes to evaluate the degree of CaMKII δ alternative splicing disorder. The data is represented as mean ± SEM. Compared with the control group* P<0.05,**P<0.01; Compared with the GSK’872+AngII group^#^ P < 0.05, ^##^P < 0.01,n=6.

### GSK’872 inhibits AngII induced necrotic apoptosis in cardiomyocytes

The study found that GSK’872 treatment significantly reduced the number of positive cells displayed by TUNEL staining and decreased the expression level of Cleaved caspase 3 in hypertrophic cardiomyocytes (P<0.05), but there was no significant difference in caspase 3 among the groups (Fig.7). These data indicate that downregulation of RIPK3 can hinder the cardiac dysfunction and necrotic apoptosis associated with myocardial hypertrophy.

**Fig. 7.**
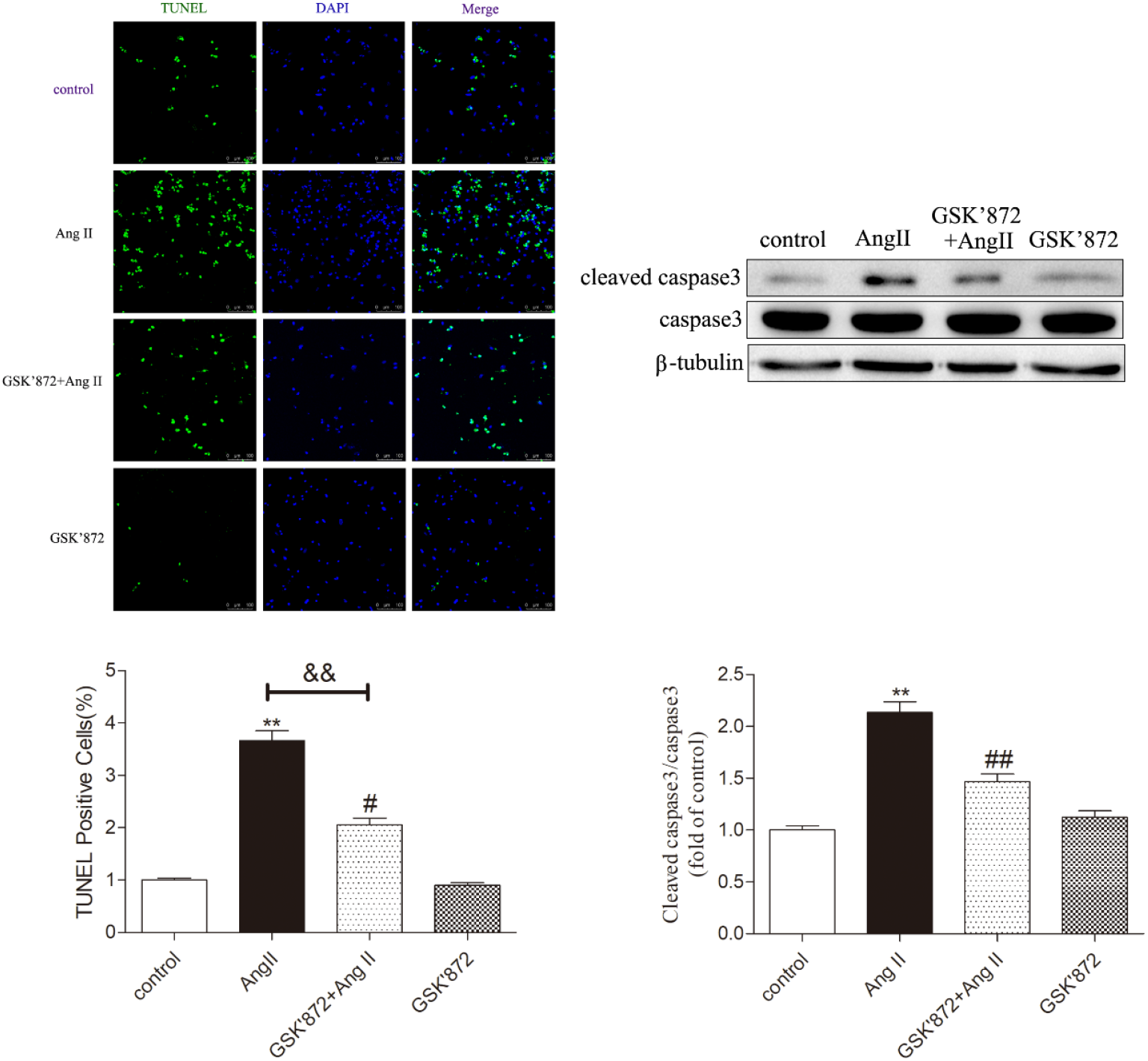
GSK’872 inhibits AngII induced necrotic apoptosis in cardiomyocytes. Using TUNEL staining to detect myocardial cell apoptosis, Bar = 50μm. Western Blot was used to detect the expression levels of caspase 3 and cleaved caspase 3 proteins to evaluate the necrotic apoptosis of myocardial cells. Significance is determined by one-way analysis of variance. The data is represented as mean ± SEM. Compared with the control group** P<0.01; Compared with the GSK’872+AngII group^##^ P < 0.01,n=6.

### GSK’872 alleviates AngII induced oxidative stress in cardiomyocytes

Our study found that under conditions of myocardial cell hypertrophy, MitoSOX staining exhibited stronger red fluorescence, while after treatment with GSK’872, its fluorescence intensity was significantly reduced, significantly lowering the levels of SOD, MDA, and T-AOC in myocardial cells (P<0.05), inhibiting oxidative stress, and enhancing the ability of myocardial cells to scavenge oxygen free radicals (Fig.8).

**Fig. 8.**
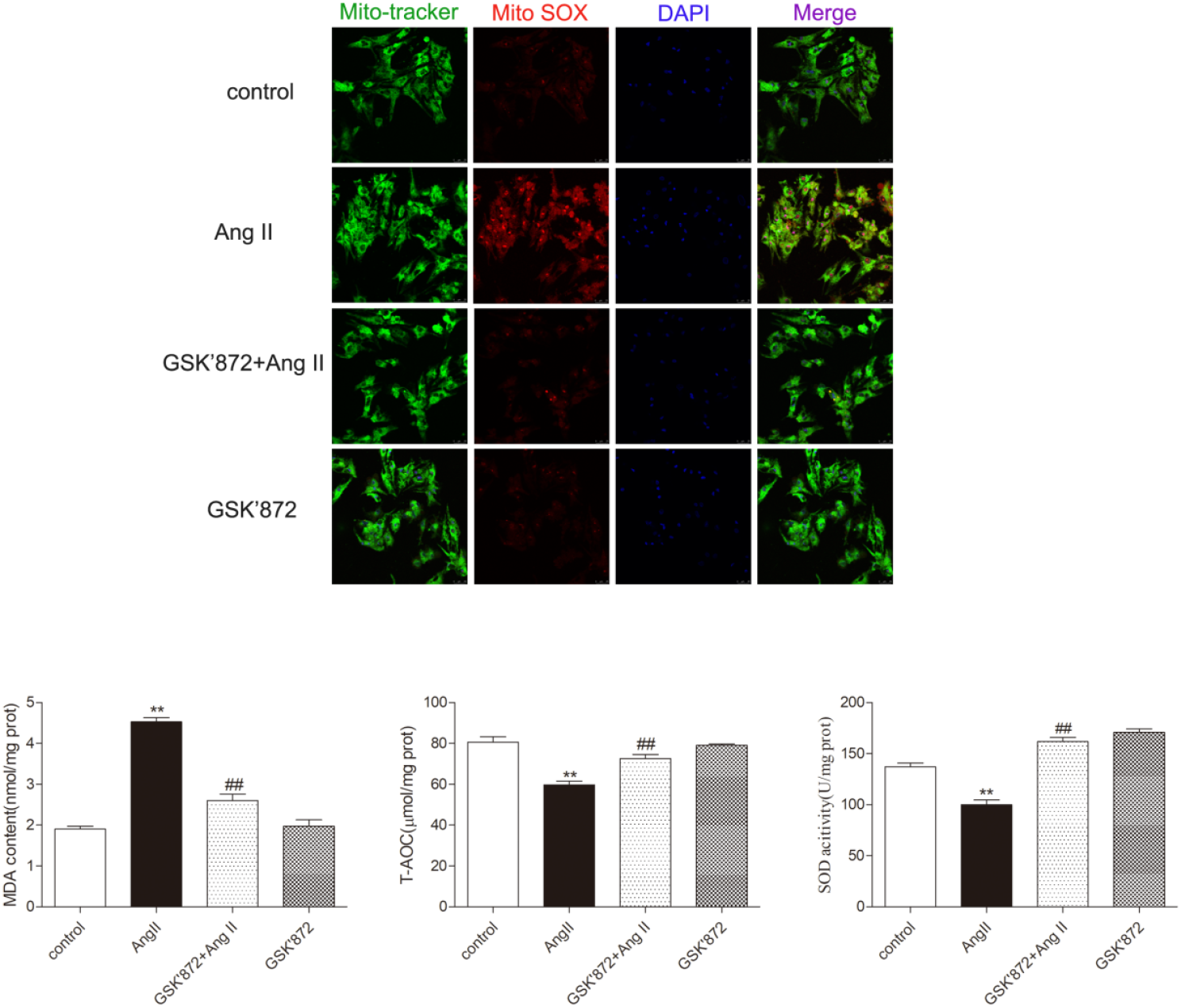
GSK’872 improves AngII induced oxidative stress and mitochondrial damage in cardiomyocytes. MitoSOX staining was used to detect intracellular superoxide in myocardial cell mitochondria, where Mito tracker was used to locate mitochondria in myocardial cells, Bar = 25μm°Measure the effects of antioxidant oxidative stress markers (SOD, MDA, T-AOC levels) in myocardial cells. The data is represented as mean ± SEM. Compared with the control group** P<0.01; Compared with the GSK’872+AngII group ^##^ P < 0.01,n=6.

### GSK’872 alleviates AngII induced mitochondrial damage and myocardial cell dysfunction in cardiomyocytes

Mitochondria, as energy producers in the heart, play a crucial role in mediating the progression of myocardial hypertrophy, and mitochondrial function can be evaluated by monitoring changes in MMP. In this study, the levels of MMP in myocardial cells were detected by JC-1 staining, as shown in Fig. 9. Compared with the control group, the red and green fluorescence ratios in the AngII group were reduced, indicating the loss of mitochondrial membrane potential (Δ ψ m) after myocardial hypertrophy. However, we further found that GSK’872 can restore the mitochondrial membrane potential of cardiomyocytes stimulated by AngII, indicating that inhibiting RIPK3 in vitro helps stabilize the MMP of mast cells and alleviate mitochondrial functional damage. Our study further found that after AngII stimulation, myocardial cells were treated with GSK’872 accompanied by a decrease in LDH levels and an increase in ATP levels (P<0.05). That is to say, downregulating RIPK3 can improve myocardial injury.

**Fig. 9.**
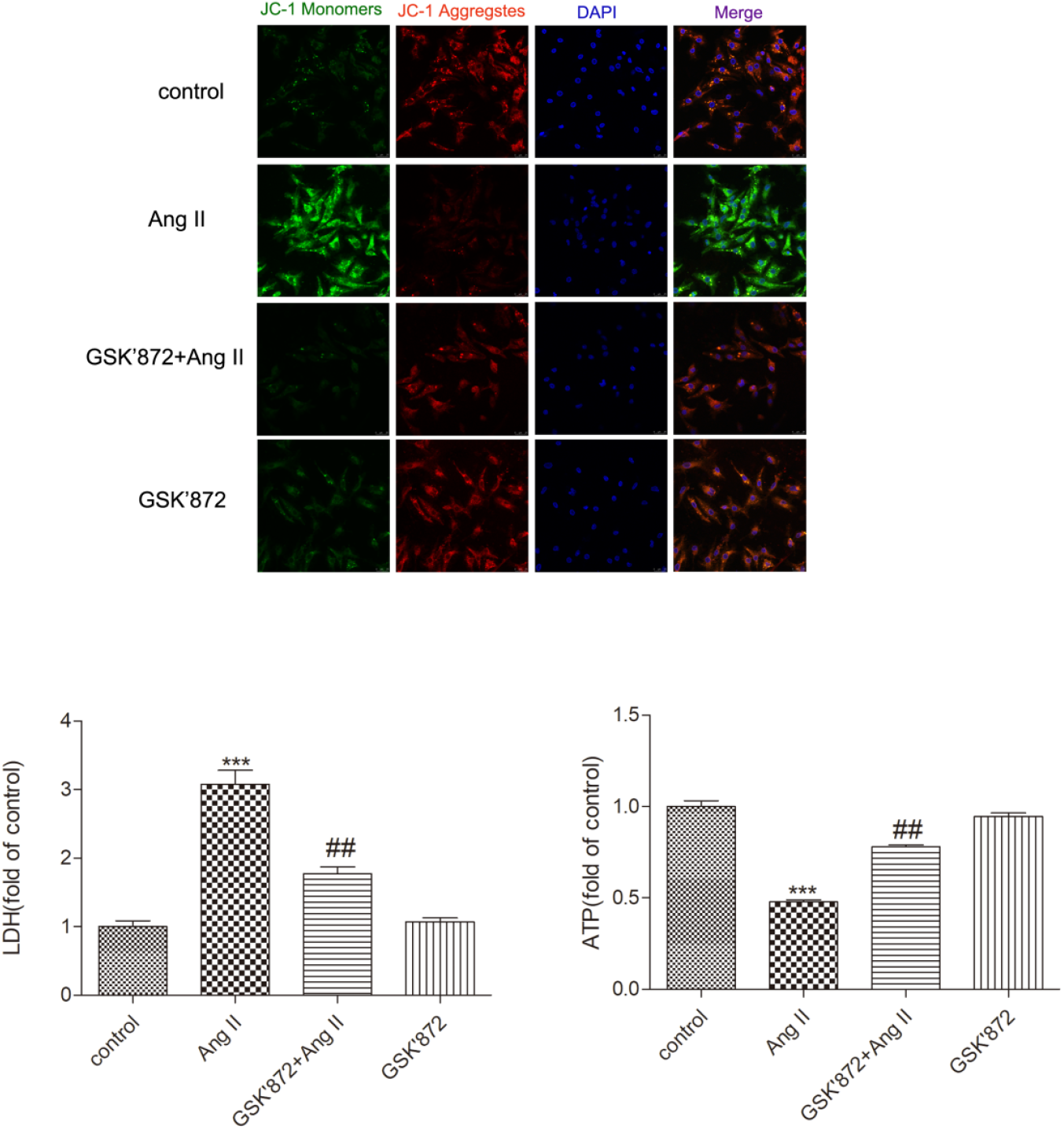
Evaluate the degree of mitochondrial damage by detecting the mitochondrial membrane potential level of myocardial cells through JC-1 staining (Bar=25 μ m). The LDH and ATP levels of myocardial cells were measured as mean ± SEM. The data is represented as mean ± SEM. Compared with the control group** P<0.01; Compared with the GSK’872+AngII group ^##^ P < 0.01,n=6.

## Discussion

The reason why myocardial hypertrophy has become a clinical medical problem of global concern is that there is currently no effective strategy to prevent and treat myocardial hypertrophy specifically. Persistent myocardial hypertrophy accumulates into dilated cardiomyopathy, which eventually develops into heart failure and sudden death^[29]^. In the pathogenesis of myocardial hypertrophy, its occurrence and development mainly involve pathological changes such as intracellular Ca^2+^overload, decreased capillary density, myocardial cell apoptosis, myocardial oxidative stress, myocardial fibrosis, and myocarditis^[30]^. These pathological changes further promote myocardial injury and dysfunction^[31-32]^. Therefore, it is particularly important to explore the regulatory effects of key target molecules on their substrates, elucidate the pathological molecular mechanisms of myocardial hypertrophy, and seek new strategies for targeted treatment of myocardial hypertrophy, targeting the pathogenic targets of myocardial hypertrophy. In this study, we used AngII to stimulate myocardial cell hypertrophy. After exposure to 100 mM AngII for 72 hours, we found a significant increase in the expression of hypertrophy genes ANP and BNP, indicating the successful establishment of a myocardial hypertrophy model. At the same time, the release of LDH in myocardial cells increased and ATP levels decreased, indicating that AngII induces myocardial cell damage. RIPK3 is a member of the RIP family and plays an important role in various cardiovascular diseases. We found that the expression of RIPK3 and p-RIPK3 increased in cardiomyocytes after AngII stimulation. However, after treatment with RIPK3 inhibitors on GSK’872, the expression and activity of RIPK3 were significantly reduced, as well as the expression of ANP and BNP. This suggests that downregulation of RIPK3 expression can alleviate AngII induced cardiomyocyte hypertrophy and injury, but its specific mechanism is still unclear.

In recent years, studies have shown a close relationship between programmed necrosis and cardiovascular disease. Patients with heart failure after myocardial infarction have elevated levels of serum tumor necrosis factor alpha, which can induce RIPK3/RIPK1 mediated programmed necrosis^[33]^. It has been reported that knockout of RIPK3 gene in atherosclerotic mice can delay the progression of atherosclerosis and alleviate the programmed necrosis of macrophages in atherosclerotic plaques, suggesting that the programmed necrosis of macrophages is a key factor in the formation of atherosclerotic plaques^[34]^. Recent studies have shown that programmed necrosis plays an important role in alcohol induced aldehyde dehydrogenase 2 gene knockout mouse cardiomyopathy^[35]^. In vitro studies have shown that palmitic acid induced programmed necrosis of myocardial cells can be reversed by inhibiting RIPK3 or RIPK1^[36]^. Previous studies have found that in mice with myocardial hypertrophy, the expression of RIPK3 in the myocardium is also significantly increased, and myocardial cell apoptosis is accelerated, inducing programmed necrosis.

The results of this study indicate that in AngII induced mouse myocardial hypertrophy tissue and AngII induced mast cells, alternative splicing significantly increases the expression of ASF and SC35, while the expression of CaMKII δ genes CaMKII δ A and CaMKII δ B significantly decreases, and the expression of CaMKII δ C significantly increases; After administration of RIPK3 inhibitor GSK’872, alternative splicing factors ASF The expression of SC35 significantly decreased, correcting or alleviating the variant disorder after CaMKII δ alternative splicing; Cultivate cardiomyocytes, GST pull down immunoprecipitation, and laser confocal fluorescence microscopy analysis showed: There is co localization between RIPK3 and ASF or RIPK3 and SC35. Tip: There is an interaction between RIPK3, ASF, and SC35; RIPK3 mediates alternative splicing of CaMKII δ by activating ASF/SC35 activity and expression levels, thereby regulating myocardial programmed necrosis. The alternative splicing variants CaMKII δ A, CaMKII δ B, and CaMKII δ C play a crucial role in the pathogenesis of myocardial hypertrophy.

It is worth noting that CaMKII is a newly discovered RIPK3 substrate. RIPK3 can induce phosphorylation of CaMKII at Thr287 and oxidation at Met281/282, triggering mPTP opening and leading to cardiomyocyte death^[37]^. Our study found that after AngII stimulation, the expression of ox CaMKII and p-CaMKII in myocardial cells increased, and the alternative splicing disorder of CaMKII δ was disrupted. Among them, the mRNA expression of CaMKII δ A and CaMKII δ B decreased, and the mRNA expression of CaMKII δ C increased. Downregulation of RIPK3 can significantly inhibit the activation of CaMKII in myocardial cells stimulated by AngII, and significantly improve the CaMKII δ alternative splicing disorder caused by AngII. These results suggest that AngII induced myocardial hypertrophy and necrotic apoptosis may be mediated by the RIPK3-CaMKII signaling axis.

Research has shown that RIPK3 enhances aerobic respiration and promotes ROS production by activating the pyruvate dehydrogenase complex. Excessive accumulation of ROS activates programmed necrosis, leading to mitochondrial dysfunction^[38-39]^.Excessive production of ROS exacerbates oxidative stress, damages mitochondria, and leads to a decrease in mitochondrial membrane potential (Δ ψ m) ^[40-41]^. The study found that MitoSOX staining was used to evaluate ROS levels in myocardial cell mitochondria; The JC-1 staining method was used to determine the Δ ψ m in the study, which is based on the penetration of mitochondrial membrane JC-1, forming aggregates that exhibit red fluorescence; In damaged cells, mitochondrial membrane depolarization occurs, and JC-1 does not accumulate in the mitochondrial matrix. JC-1 only forms monomers that appear fluorescent green; RIPK3 knockout in diabetes cardiomyopathy mice can significantly improve the ultrastructure of myocardial mitochondria^[19,42]^. The results of this study indicate that: After AngII induced mouse myocardial hypertrophy, transcriptome analysis revealed downregulation of genes related to mitochondrial energy metabolism, such as acyl coenzyme metabolism and fatty acid beta oxidation. Transmission electron microscopy revealed disordered arrangement, swelling, and cristae rupture of cardiac mitochondria; After knocking out the RIPK3 gene or administering the RIPK3 inhibitor GSK’872, mitochondrial abnormalities improved. In AngII induced mast cells, mitochondrial membrane potential was significantly reduced, and after administration of RIPK3 inhibitor GSK’872, mitochondrial membrane potential was significantly increased. Tip: RIPK3 mediates CaMKII δ alternative splicing to regulate mitochondrial function; Improving mitochondrial function, inhibiting ROS levels, reducing cellular oxidative stress, increasing mitochondrial membrane potential, protecting mitochondria from depolarization damage, thereby reducing myocardial injury.

In summary, RIPK3 mediates alternative splicing of CaMKII δ, and activation of CaMKII δ causes disruption of alternative splicing of CaMKII δ variants, leading to programmed necrosis of myocardial cells; RIPK3 inhibitors can decrease the activity of CaMKII, Correction of CaMKII δ alternative splicing disorder, alleviation of oxidative stress, and reduction of programmed necrosis. The inhibitor GSK’872 of the target molecule RIPK3 has the potential to become a new targeted drug for the treatment of AngII induced myocardial hypertrophy, providing a molecular basis and new clinical treatment strategies for targeted prevention and treatment of myocardial hypertrophy, dilated cardiomyopathy, and heart failure.

## Materials and Methods

### 1. Primary cardiomyocyte culture

Take 1-3 day old SD neonatal rats, disinfect the skin on the chest and abdomen, and perform thoracic incision surgery to remove the heart. Immediately wash with pre cooled phosphate buffered saline containing 1% antibiotic (penicillin streptomycin). Subsequently, the heart was cut into pieces and digested in 0.25% trypsin (Beyotime, Shanghai, China). Gently shake in a preheated 37 °C water bath, discard the supernatant after the first digestion for 5 minutes, and then collect the supernatant every 3 minutes. Repeat the above steps about ten times until complete digestion. After inhaling the supernatant, filter the cells into a 100ml beaker and centrifuge at 1200 rpm for 5 minutes. Resuspend the centrifuged cell pellet in DMEM containing 10% fetal bovine serum (FBS, GIBCO, Carlsbad, CA, USA). Cultivate the cell suspension in a 37 °C incubator. To remove cardiac fibroblasts, transfer the unattached cells to fresh culture medium after 4 hours for subsequent cultivation.

### 2. GSK’872 treatment

Before pretreatment with RIPK3 inhibitor GSK’872, myocardial cells need to be starved in FBS free medium for 10 hours when the fusion rate of myocardial cells reaches around 80%. Subsequently, cardiomyocytes were treated with culture medium containing GSK’872 (10 μ M, GlpBio, USA) and 10% fetal bovine serum for 4 hours. Afterwards, the culture medium was changed to 10% fetal bovine serum medium containing AngII (100 μ M. Sigma) or without AngII.

### 3. Western blot analysis

Proteins were extracted from myocardial cells and separated by sodium dodecyl sulfate polyacrylamide gel electrophoresis (SDS-PAGE). Then transfer the protein onto a polyvinylidene fluoride (PVDF) membrane (Billerica Millipore, Massachusetts, USA). Seal the membrane with TBST buffer containing 5% skim milk at room temperature for 2 hours, then incubate overnight with relevant primary antibodies at 4°C. The main antibodies used are as follows: anti-RIPK3 (1:1000, Novus); Anti ANP, anti BNP, anti p-RIPK3, anti ASF, anti SC35, and anti CaMKII (1:1000, abacm); Anti-RIPK1, anti-p-MLKL, anti-caspase-3, and anti cleavage caspase-3 (1:1000, cell signaling technology); Anti ox CaMKII (1:1000, Millipore, Germany); Anti-p-CaMKII (1:500, Santa Cruz Biotechnology); Anti GAPDH (1:5000, Shanghai Kangchen Biotechnology Co., Ltd.); Anti beta microtubule protein (1:5000, CMCTAG). Subsequently, the membrane was cultured at room temperature for 2 hours using appropriate horseradish peroxidase (HRP) as a secondary antibody (1:5000, Jinqiao, Zhongshan, Beijing, China). Using a bio-radchemidocXRS+imaging system (Thermo Fisher Scientific, Rockford, Illinois, USA), protein bands were observed through enhanced chemiluminescence (ECL). Quantitative analysis was conducted using ImageJ software.

### 4. Quantitative real-time RT-PCR

Accordingtothemanufacturer’splan,total RNA was extracted from myocardial cells using theHiScript®IIQRT SuperMix for qPCRkit (vazyme, Nanjing, China). Quantify and balance the total RNA before using AceQ®The Universal SYBR qPCR Master Mix kit (Nanjing Vazyme, China) was used to reverse transcribe it into cDNA. Perform three real-time fluorescence quantitative PCR analyses on the cDNA of each group of myocardial cells, with 18S as the reference gene cluster. The table reports the sequences of all primers.

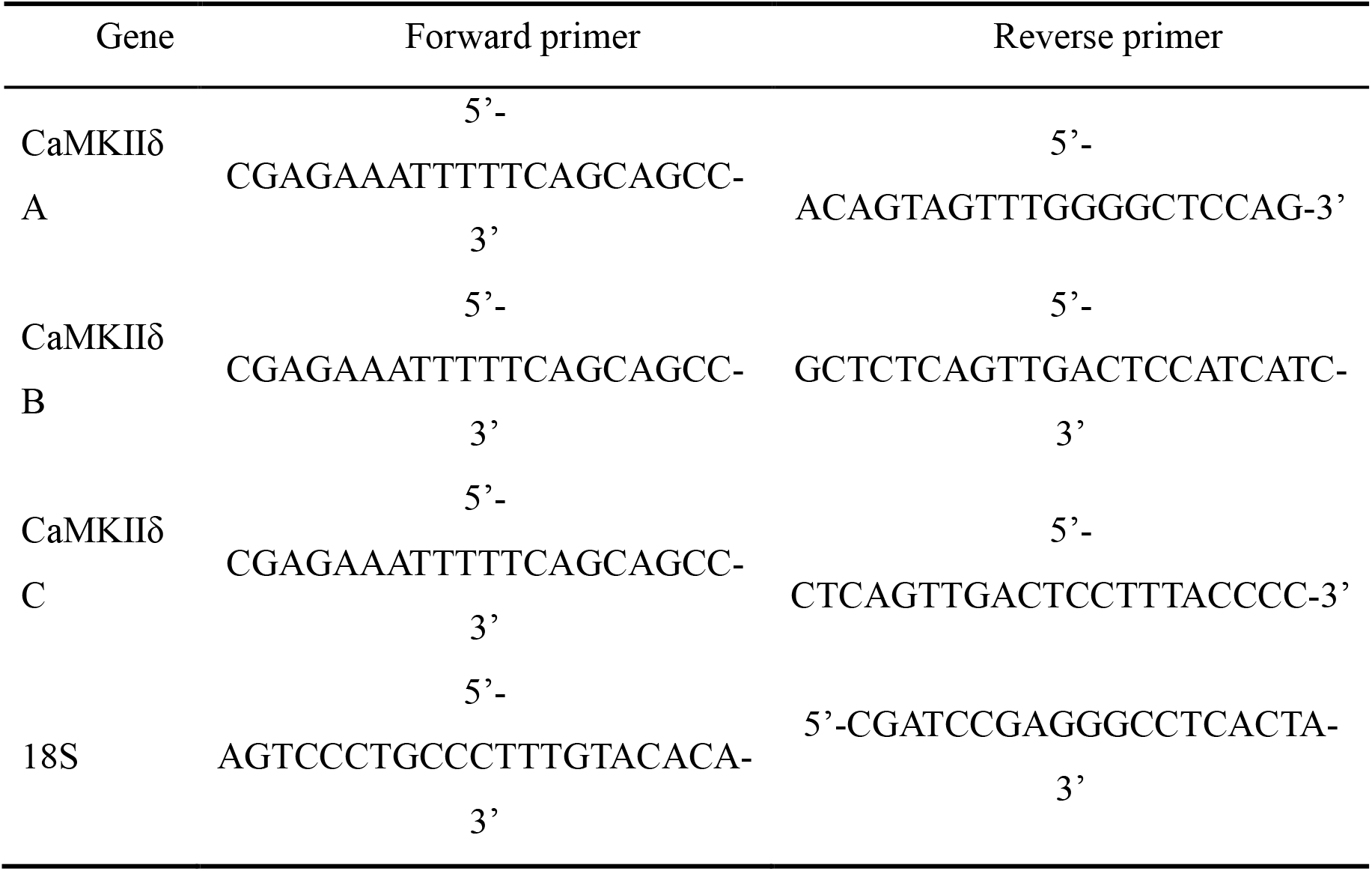

### 3.4. In situ terminal transferase labeling (TUNEL method)

According to the one-step TUNEL apoptosis detection kit (Biyun Tian Company, China), apoptotic cardiomyocytes were detected under the catalysis of TDT, and the addition of fluorescein dUTP resulted in green fluorescence. The nuclei were labeled with DAPI and showed blue fluorescence. Fluorescence changes were observed using a laser confocal microscope (LEICA Company, Germany).

### 5. Mitochondrial Superoxide Assessment

Inorder to evaluate the oxidative stress level of myocardial cells, MitoSOX targeted transmembrane fluorescent staining can be used to detect its level. MitoSOX Red (diluted with PBS at a ratio of 1:1000) and Mito Tracker Green staining solution (diluted with PBS at a ratio of 1:5000) are prepared and shaken evenly. The mixture is preheated in a 37 °C constant temperature incubator, andMitoSOX Red and Mito Tracker Green co staining solutions are prepared to stain myocardial cells. The cells are incubated in a 37 °C constant temperature incubator in the dark for 15 minutes.Afterwards, DAPI stained cell nuclei were added, and an appropriate amount of anti fluorescence quencher was dropped onto the glass slide. The slide was then inverted and observed under a confocal microscope.

### 6. Mitochondrial membrane potential (JC-1) detection

In order to evaluate the mitochondrial membrane potential (Δψm) of cardiomyocytes, following the instructions of the JC-1 kit (Beyotime, Shanghai, China), cardiomyocytes were stained with preheated JC-1 staining solution for 20 minutes at 37 °C in the dark, and then stained with JC-1 staining buffer (1×). Next, add DAPI staining to the cell nucleus. Finally, observe the myocardial cells using a confocal laser microscope and take photos.

### 7. Determination of LDH, ATP, SOD, MDA, and T-AOC in cells

According to LDH cytotoxicity assay kit and CellTiterLumi™ instructions for the vitality detection kit (Biyuntian, Shanghai, China) are to detect the LDH content in cell culture medium, evaluate the damage of myocardial cells after AngII stimulation, and assess the ATP level in myocardial cells. Using a microplate reader (BioTek, USA), perform absorbance detection at 490nm to calculate cell toxicity and chemiluminescence detection to calculate cell viability. In order to determine the effect of antioxidant stress markers in cardiomyocytes, cardiomyocytes were seeded in 6-well plates and cultured at 37 ° C. After the cells grew to 80% fusion, they were processed according to experimental needs. The activity or content of superoxide dismutase (SOD), total antioxidant capacity (T-AOC), and malondialdehyde (MDA) in cardiomyocytes were measured using a commercial kit (Nanjing Jiancheng Bioengineering Institute) according to the instructions.

### 8. Statistical analysis

The experimental data are presented as mean ± standard error (± S.E.M) indicates that GraphPad Prism software was used to process the experimental data, and one-way ANOVA and Student Newman Keuls (SNK) were used to test the statistical analysis of the data. When P<0.05, the experimental data is considered statistically significant.

## Author Contributions

J.Z. were involved in the conception and design of the work, analysis of data, and manuscript drafting. H Y.Y Y and A L.W. were involved in the acquisition and analysis of data. P W. were involved in the design of the research report and proofreading of the original manuscript. All authors have read and agreed to the published version of the manuscript.

